# Thalamic nucleus reuniens coordinates prefrontal-hippocampal synchrony to suppress extinguished fear

**DOI:** 10.1101/2022.11.11.516165

**Authors:** Michael S. Totty, Karthik R. Ramanathan, Jingji Jin, Shaun E. Peters, Stephen Maren

**Affiliations:** Department of Psychological and Brain Sciences, Texas A&M University, College Station, Texas; Institute for Neuroscience, Texas A&M University, College Station, Texas

## Abstract

Traumatic events result in vivid and enduring fear memories. Suppressing the retrieval of these memories is central to behavioral therapies for pathological fear. The medial prefrontal cortex (mPFC) and hippocampus (HPC) have been implicated in retrieval suppression, but how mPFC-HPC activity is coordinated during extinction retrieval is unclear. Here we show that after extinction training, coherent theta oscillations (6-9 Hz) in the HPC and mPFC are correlated with the suppression of conditioned freezing in male and female rats. Inactivation of the nucleus reuniens (RE), a thalamic hub interconnecting the mPFC and HPC, reduces extinction-related Fos expression in both the mPFC and HPC, dampens mPFC-HPC theta coherence, and impairs extinction retrieval. Conversely, theta-paced optogenetic stimulation of RE augments fear suppression and reduces relapse of extinguished fear. Collectively, these results demonstrate a novel role for RE in coordinating mPFC-HPC interactions to suppress fear memories after extinction.

## Introduction

For many individuals, traumatic events result in vivid and enduring memories that drive unrelenting and pathological fear^1^. Behavioral interventions, such as prolonged exposure therapy, rely upon extinction learning to suppress emotional memories and dampen fear responses^2,3^. Considerable work demonstrates that the mPFC is critical for extinction learning and may be an essential neural substrate for the suppression of episodic fear memories after extinction^4–7^. Work in both humans and animals suggests that interactions between the mPFC and HPC are critical for the suppression of episodic memories^8,9^. Indeed, inhibiting the retrieval of context-inappropriate memories, a hallmark of retrieval suppression, is critical for the expression of context-dependent extinction^10,11^. Hence, mPFC-HPC interactions may prevent intrusive fear memories from disrupting extinction retrieval^12–14^, though the mechanisms supporting this function are unknown.

Neural oscillations facilitate interactions between brain regions by coordinating and synchronizing neural activity^15^. Investigations into the neural oscillations underlying aversive learning and memory^16^ have discovered that theta-range oscillations act to synchronize limbic structures during the retrieval of both fear^16–21^ and extinction memories^18,22,23^. For example, considerable evidence implicates synchronization of prefrontal-hippocampal oscillations in memory retrieval^24–28^. However, it is unclear how the mPFC influences the retrieval of hippocampus-dependent fear and extinction memories insofar as the mPFC has only sparse projections to the HPC^29,30^. One candidate brain region coordinating interconnecting the mPFC and HPC is the thalamic nucleus reuniens (RE). The reuniens is a small structure in the ventral midline thalamus that bidirectionally connects the mPFC and HPC^31^ and is a critical hub for coordinating mPFC-to-HPC interactions^32^. Indeed, Ramanathan and colleagues found that the RE and mPFC➝RE projections, are necessary for the encoding and retrieval of both auditory and contextual extinction memories^33,34^. Inactivation of the RE also impairs the precision of contextual fear memories^34,35^ a process that requires both the HPC and mPFC ^36,37^. Additionally, inactivating the RE has been reported to impair mPFC-HPC synchrony^38–40^. Collectively, these data suggest that RE may be critical for mPFC-HPC synchrony during the retrieval of extinction memories.

Here, we test this specific hypothesis using pharmacological, optogenetic, and electrophysiological methods in behaving rats. We found that the mPFC and RE display distinct 3-6 Hz and 6-9 Hz oscillations that are positively and negatively, respectively, correlated to freezing behavior during extinction acquisition and retrieval. Extinction retrieval was associated with increased theta coherence between the mPFC and HPC. Pharmacological inactivation of the RE induced fear relapse after extinction, impaired c-Fos immediate early gene expression in the mPFC and HPC, and abolished mPFC-HPC theta synchrony. Non-conditioned controls demonstrate that increases in freezing behavior induced by RE inactivation are specific to memory retrieval. Finally, we show that theta-paced optogenetic stimulation of the RE is sufficient to block fear relapse. These data demonstrate that RE is a critical hub for the retrieval of safe extinction memories and may be a novel therapeutic target for the suppression of intrusive traumatic memories.

## Results

### Fear extinction is characterized by mPFC-HPC theta coherence

To examine the role of mPFC-HPC theta synchronization in fear extinction, we simultaneously recorded local field potentials (LFPs) (Fig. 1A) from both the prelimbic (PL) and infralimbic (IL) regions of the mPFC and the CA1 region of the dorsal HPC (Fig. 1C) during extinction acquisition and retrieval. Rats (*n* = 6) were conditioned using a standard auditory fear conditioning procedure (Fig. 1B). After conditioning, recordings were made during the context exposure, fear extinction, and extinction recall sessions.

**Figure 1.**
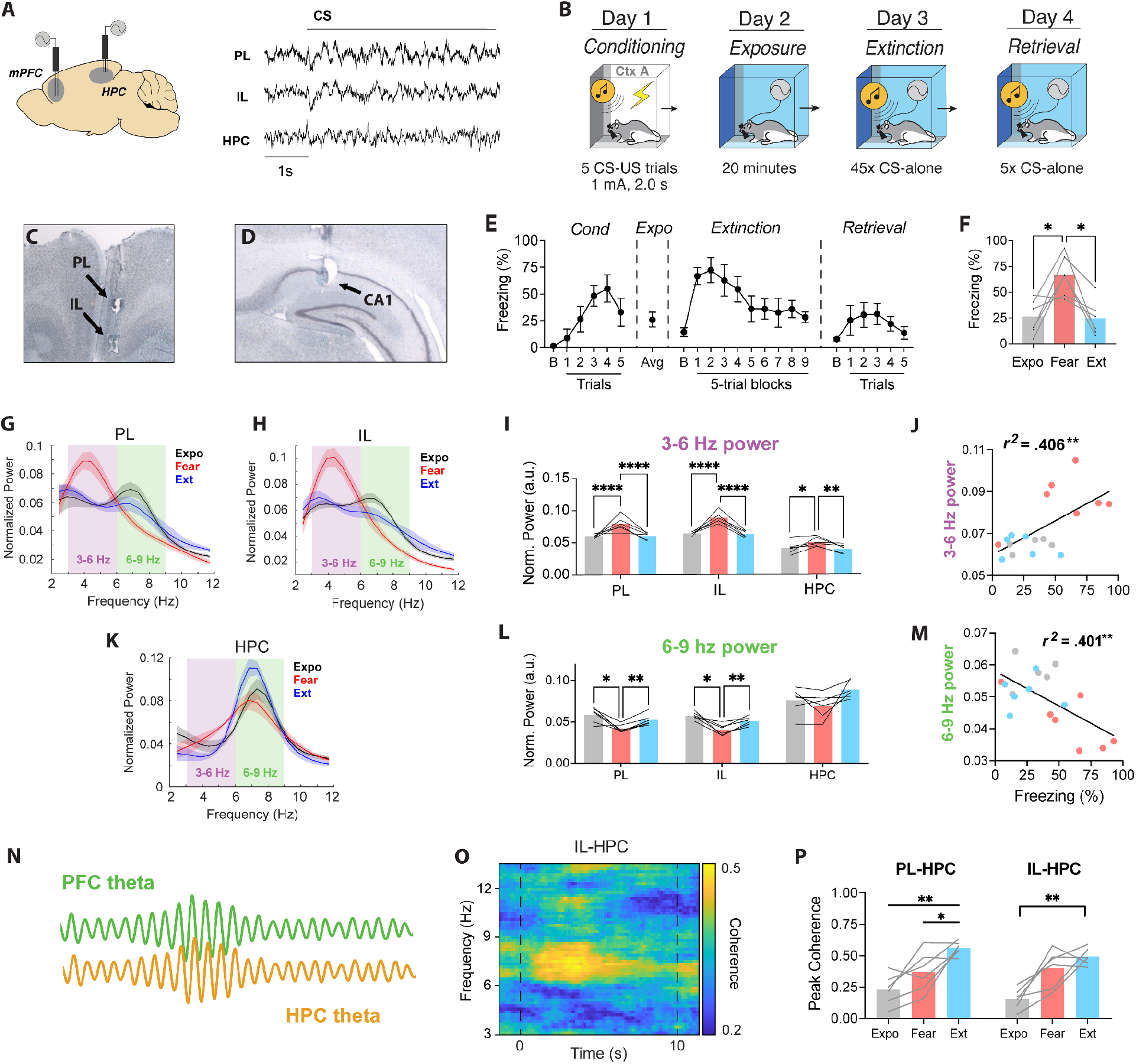
In vivo electrophysiological recordings during context exposure, fear extinction, and extinction retrieval. (A) Illustration and representative traces of mPFC and HPC local field potential recordings (LFPs). (B) Behavioral timeline of the fear conditioning protocol. Representative photomicrographs show mPFC (C) and HPC (D) electrode placements. (E) Averaged freezing behavior (*n* = 6) during fear conditioning, context exposure, extinction, and extinction retrieval. (F) Average freezing during context exposure (Expo), fear retrieval during the first five trials of extinction (Fear), and the five extinction retrieval trials (Ext). (G, H, K) Spectral power of theta-range oscillations in the prelimbic (PL) and infralimbic (IL) cortex and the HPC during 10 sec CS presentations across days. (I, L) Average theta-range power across behavioral time points. (J,M) Linear regression of freezing vs IL theta-range power during Expo, Fear and Ext. (N) Cartoon illustrating synchronous theta oscillations. (O) Example coherence plot depicting high mPFC-HPC theta coherence during extinction retrieval. (P) Average peak coherence across behavioral points of the interest. All data are means ± s.e.m.s; \**p* < 0.05; \*\**p <* 0.01; \*\*\**p <* 0.001; \*\*\*\**p <* 0.0001.

As shown in Fig. 1E, all animals acquired fear of the CS, which was manifest as increased freezing across conditioning trials (main effect of trials: *F*_*5*,*20*_ = 5.44, *p* = .003). The following day, all rats were exposed to the extinction context f-or 20 minutes where they displayed low levels of freezing. During extinction training, all rats showed high fear to the CS presentations during the early extinction trials and freezing decreased during successive CS presentations (main effect of trials: *F*_*9*,*36*_ = 6.053, *p* < .0001). CS-elicited freezing behavior remained low during retrieval testing the following day, demonstrating a robust extinction memory.

We chose to focus our electrophysiological analyses on three sessions in which animals showed different levels of conditioned fear: context exposure (‘Expo’, low fear), early extinction (‘Fear’, high fear; first 5 trials of extinction), and extinction retrieval (‘Ext’, suppressed fear; first 5 trials of extinction retrieval). As shown in in Fig 1F, conditioned freezing during these sessions was significantly higher early in the extinction session (‘Fear’) relative to the exposure (‘Expo’) and extinction retrieval (‘Ext’) sessions (main effect of Days: *F*_*2*,*10*_ = 9.664, *p* = .006). Examining the LFP recordings, we found that the expression of conditioned freezing during the earliest extinction trials (‘Fear’) was associated with greater 3-6 Hz power (main effect of Day: *F*_*2*,*30*_ = 46.53, *p* < .0001) in the IL (Fear vs Ext post hoc: *p* < .0001), PL (Fear vs Ext post hoc: *p* < .0001), and HPC (Fear vs Ext post hoc: *p* = .0062) relative to the extinction retrieval session (Fig. 1G-M). In contrast, the suppression of conditioned freezing during extinction extinction retrieval (‘Ext’) was associated with greater 6-9 Hz power (main effect of Day: *F*_*2*,*30*_ = 13.89, *p* < .0001) in the IL (Fear vs Ext post hoc: *p* = .002) and PL (Fear vs Ext post hoc: *p* = .0092), but not the HPC (Fear vs Ext post hoc: *p* = .128) relative to the early extinction trials. Moreover, we found that, across all sessions, 3-6 Hz power was positively correlated with conditioned freezing in both IL (*r*^*2*^ = .406, *p* = .004) and PL (*r*^*2*^ = .235, *p* = .042) (Fig. 1J), whereas freezing behavior was negatively correlated with 6-9 Hz power in the IL (*r*^*2*^ = .401, *p* = .005) (Fig. 1M). The correlation between freezing behavior and 6-9 Hz oscillations in the PL was also negative, but was not statistically significant (*r*^*2*^ = .207, *p* = .058). This further demonstrates that freezing behavior is associated with an increase in ∼4 Hz oscillatory power in the mPFC.

Next, we sought to determine if the retrieval of extinction memories was associated with increased mPFC-HPC theta synchrony (Fig. 1N). Considering that 6-9 Hz oscillations are negatively correlated with conditioned freezing (e.g., Fig 1M), we predicted that fear extinction would also be associated with mPFC-HPC coherence in this frequency band. Indeed, we found that mPFC-HPC coherence is primarily centered at ∼8 Hz (Fig. 1O). The HPC showed increased coherence (6-9 Hz) with the IL (main effect: *F*_*2*,*10*_ = 16.69, *p* = .0007; Expo vs Ext post hoc: *p* = .0006) and PL (main effect: *F*_*2*,*10*_ = 13.94, *p* = .0013; Expo vs Ext post hoc: *p* = .001) during extinction retrieval compared to context exposure (Fig. 1P). This demonstrates that the retrieval of extinction memories is associated with 6-9 Hz theta synchrony between the mPFC and HPC.

### The thalamic nucleus reuniens exhibits theta-range oscillations during extinction that correlate with conditioned freezing

We next sought to determine if the RE also displays similar oscillations across fear extinction (Fig. 2A). To this end, a multielectrode array was implanted in the RE one week prior to behavioral testing; bipolar LFP recordings were conducted by referencing one channel off another intra-RE channel to remove volume conducted signals from external sources (Fig. 2B). All rats (*n* = 4) were fear conditioned as previously described.

**Figure 2.**
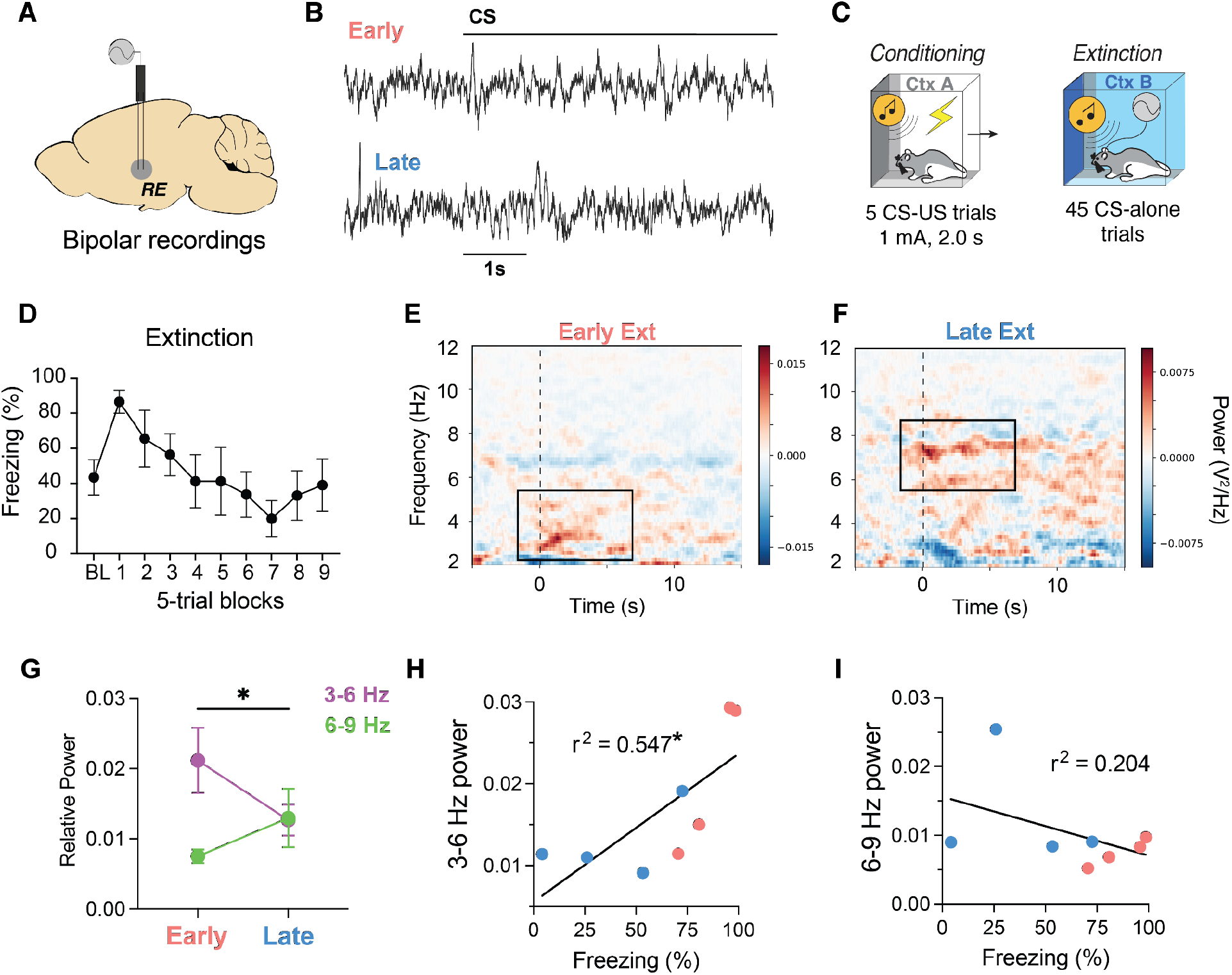
The nucleus reuniens displays theta-range rhythms that correlate with conditioned freezing across extinction learning. (A) Illustration and (B) raw traces of bipolar electrophysiological field recordings from the RE. (C) Behavioral timeline showing that field recordings took place during fear extinction after all rats were conditioned the previous day. (D) Average freezing behavior (*n* = 4) during fear extinction training.. Baseline subtracted spectrograms showing increased 3-6 Hz power (E) during the first five trials of extinction (Early) and increased 6-9 Hz power (F) during the last five trials of extinction (Late). (G) Average relative power of 3-6 and 6-9 Hz power during Early compared to Late extinction. Linear regression of freezing behavior vs 3-6 Hz power (H) and 6-9 Hz power (I). All data are means ± s.e.m.s; \**p* < 0.05.

During fear extinction, rats exhibited high levels of conditioned freezing behavior that extinguished over 45 CS-alone trials (Fig. 2C-D). Like the previous experiment, we chose to restrict our electrophysiological analyses to the first and last 5 trials of extinction to examine differences in high and low fear states, respectively. Similar to the mPFC and HPC, we observed prominent 3-6 Hz oscillations (Fig. 2E) during the first five trials of extinction (Early Ext) and this switched to 6-9 Hz oscillations (Fig. 2F) during the last five trials (Late Ext). This was confirmed by focusing on the relative spectral power during the CS (Fig. 2G), showing that 3-6 Hz power decreased, whereas 6-9 Hz power increased, from Early to Late Ext (Frequency X Block interaction: *F*_*1*,*6*_ = 7.376, *p* = .035). We further show that 3-6 Hz rhythms positively correlated with freezing behavior (*F*_*1*,*6*_ = 7.233, *r*^*2*^ = 0.547, *p* = .036), whereas 6-9 Hz rhythms displayed a significantly different negative association with freezing behavior (*F*_*1*,*12*_ = 7.64, *p* = .017). These data confirm that the RE displays similar rhythms as the PFC and HPC across fear extinction.

### Fear relapse induced by RE inactivation impairs c-Fos expression in the mPFC and HPC

Previous work found that RE is critical for the retrieval of extinction memories (refs), and it is likely that the RE mediates this process by driving neural activity in the mPFC and HPC. To test this hypothesis, we first sought to determine if fear relapse induced by RE inactivation impaired immediate-early gene (c-Fos) expression, a proxy for neural activity, in the mPFC and HPC. To this end, rats were implanted with a single midline cannula targeting the RE. One week after surgery, all rats underwent standard fear conditioning and extinction. Conditioning and extinction behavior was similar to that observed in the previous experiments and there were no differences between drug groups (*p* > 0.3). As shown in Fig 3A, rats that were infused with muscimol (MUS; *n* = 14) exhibited poor extinction recall compared to saline (SAL; *n* = 15) controls (main effect of Drug: *F*_*1*,*27*_ = 8.66, *p* = .007). All rats were then sacrificed 90 minutes after retrieval testing to examine Fos expression. We found that MUS animals displayed decreased Fos expression, an indication of reduced neural activity, in the HPC (Fig. 3C; *t*_*27*_ = 3.26, *p* = .003) as well as both PL (*t*_*27*_ =2.59, *p* = .015) and IL (*t*_*27*_ = 2.45 *p* = .021) subregions of the mPFC (Fig. 3D), compared to SAL controls. Importantly, Fos expression in the medial geniculate nucleus (MGN), a thalamic auditory region, was not affected by RE inactivation (*t*_*27*_ = 0.71, *p* = .485). These data suggest that fear relapse induced by RE inactivation is associated with reduced neuronal activity in the prefrontal-hippocampal network.

**Figure 3.**
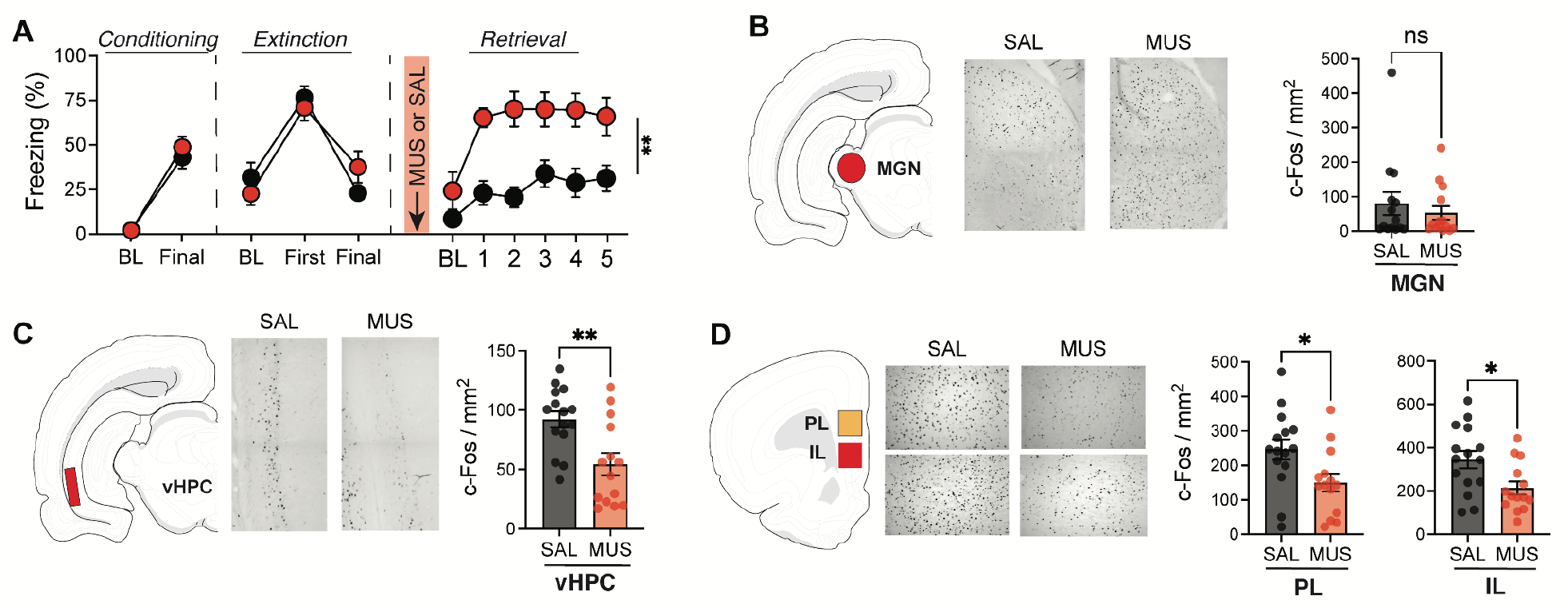
Pharmacological inactivation of RE suppreses Fos expression in mPFC and HPC during after relapse of extinguished fear. (A) Freezing behavior during conditioning, extinction, and extinction retrieval after saline (SAL; *n* = 15) or muscimol (MUS; *n* = 14) infusions into the RE. Fear relapse induced by MUS inactivation of the RE did not impair c-Fos expression in the MGN (B) but did impair c-Fos expression in both the vHPC (C) and both IL and PL subregions of the mPFC (D). All data are means ± s.e.m.s; \**p* < 0.05; \*\**p <* 0.01.

### Pharmacological inactivation of RE impairs both mPFC-HPC synchrony and extinction retrieval

It has previously been shown that RE is critical for both the retrieval of extinction memories^33,34^ and mPFC-HPC synchrony during a working memory task^38^. However, it is unknown if inactivating the RE impairs mPFC-HPC synchrony associated with fear extinction. To test this hypothesis, we implanted electrodes in the mPFC and HPC and a cannula in the RE to record mPFC and HPC LFPs while pharmacologically inactivating the RE (Fig. 4A). One week after surgery, all animals (*n* = 11) underwent standard fear conditioning and extinction as previously described (Fig. 4B). For extinction retrieval testing, rats were either infused with either the GABA_A_ agonist, muscimol (MUS), or saline (SAL) immediately prior to testing (drug infusions were counterbalanced for test order).

**Figure 4.**
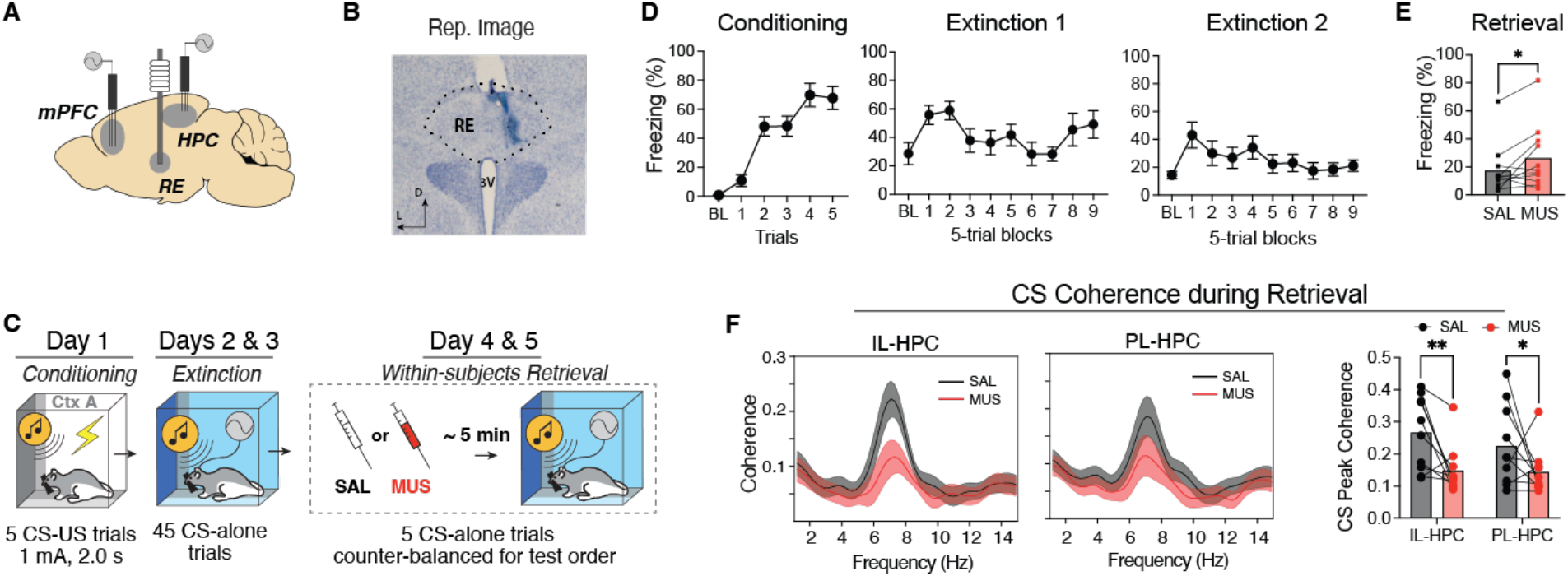
Pharmacological inactivation of RE impairs mPFC-HPC synchrony. (A) Illustration depicting electrode and cannula placements. (B) Representative photomicrograph showing injector tip placement and muscimol spread in the RE. (C) Behavioral timeline. (D) Average freezing behavior (*n* = 11) across fear conditioning and extinction. (E) Average freezing behavior showing that muscimol inactivation of the RE impairs extinction retrieval. (F) Average theta-range spectral coherence between the mPFC and HPC after either muscimol or saline infusion in the RE. All data are means ± s.e.m.s; \**p* < 0.05; \*\**p <* 0.01.

As shown in Figure 4D, all animals acquired conditioned freezing to the CS (main effect of trials: *F*_*5*,*45*_ = 26.60, *p*< .0001) and subsequently extinguished freezing to the CS (main effect of blocks: *F*_*2*,*81*_ *= 2*.*638, p =* .0099). During retrieval testing (Fig. 4E), muscimol inactivation of the RE increased freezing (*t*_*10*_ = 2.269, *p* = .0467), effectively impairing extinction retrieval as we have previously reported^33,34^. To determine if RE inactivation impaired mPFC-HPC coherence, coherence was averaged across the 5 retrieval trials for each animal and the peak coherence value in 6-12 Hz range was found (Fig. 4F). Comparing these peak coherence values, we found that muscimol inactivation of the RE impaired both PL-HPC (muscimol vs saline post hoc: *p* = .0402) and IL-HPC (muscimol vs saline post hoc: *p* = .004) 6-9 Hz theta coherence (main effect of drug: *F*_*1*,*18*,_ = 5.976, *p* = .037). This demonstrates that RE is indeed critical to both the retrieval of extinction memories and its associated mPFC-HPC theta synchrony.

### Optogenetic inhibition of RE selectively impairs extinction memory retrieval

Pharmacological inhibition of the RE may have disrupted performance during retrieval testing either by impairing contextual processing prior to CS onset or by affecting extinction memory retrieval during presentation of the extinguished CS. To assess whether RE is involved in extinction retrieval, we adopted an optogenetic strategy to silence the RE immediately prior to CS delivery, leaving the RE functionally intact during the pre-CS baseline period (when animals sample the context). We also included a non-conditioned control to determine whether increases in freezing associated with RE inactivation are due to nonspecific increases in anxiety^41^. To this end, rats were first injected with an adeno-associated virus encoding with a red-light activated, inhibitory opsin (AAV8-CaMKII-Jaws-GFP) or a control fluorescent protein (AAV8-CaMKII-GFP) into the RE, along with an optic fiber, four weeks prior to behavioral testing (Fig. 5A). Half of the rats expressing the Jaws virus were conditioned (Jaws; *n* = 12) while the other half were merely exposed to the CS without US reinforcement to serve as a non-conditioned control group (Control; *n* = 8). All rats expressing the control GFP virus were conditioned (GFP; *n* = 10).

**Figure 5.**
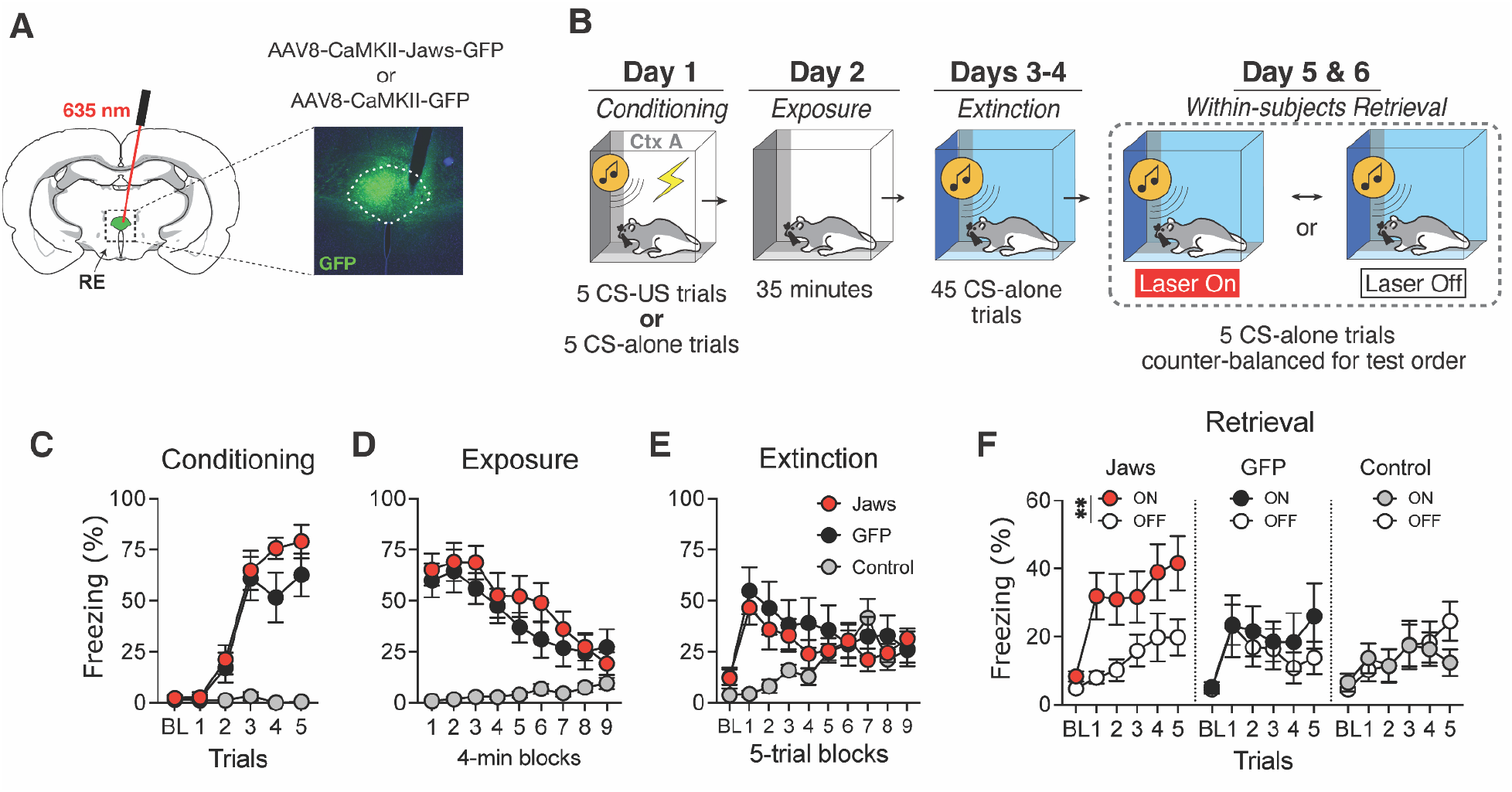
Optogenetic inactivation of the RE selectively impairs extinction memory retrieval. (A) Illustration depicting fiber placement and intracranial injections of either the active Jaws (AAV8-CaMKII-Jaws-GFP) or GFP control virus (AAV8-CaMKII-GFP) into the RE. (B) Experimental timeline. Average freezing behavior during fear conditioning (D), context exposure (E), and extinction (F). (G) Average freezing behavior during extinction retrieval showing that optogenetic inhibition of the RE selectively increases freezing in conditioned animals expressing the active Jaws virus (*n* = 13), but not GFP (*n* = 10) or non-conditioned control animals (*n* = 8). All data are means ± s.e.m.s; \**p* < 0.05; \*\**p <* 0.01.

During conditioning (Fig. 5D), both Jaws and GFP animals acquired high levels of conditioned freezing, whereas the non-conditioned Control group exhibited low levels of freezing behavior (Trial x Group interaction: *F*_*10, 135*_ = 9.067, *p* < .0001). The following two days, all rats were first extinguished to the conditioning context (Fig. 5E; Block x Group interaction: *F*_*16, 216*_ = 5.024, *p* < .0001) and then to the auditory CS in a novel context (Fig. 5F; Block x Group interaction: *F*_*18, 234*_ = 3.629, *p* < .0001). For retrieval testing, all rats were tested for extinction memory either with (ON) or without (OFF) constant red laser stimulation using a counterbalanced, within-subject design. If RE inactivation produces an anxiogenic state, one would expect both the Jaws and Control groups to show increased fear during retrieval testing. However, planned comparisons revealed optogenetic inactivation of the RE only increased freezing in the Jaws animals that had been conditioned (Laser x Group interaction: *F*_*2, 28*_ = 3.095, *p* = .061; ON vs OFF post hoc: *p* = .005), but not the GFP (*p* = .811) or Control animals (*p* = .989) (Fig. 5G-H). This effectively demonstrates that freezing induced by RE inactivation is in fact specific to the relapse of extinguished fear, rather than a non-specific anxiogenic effect. Moreover, these results reveal that RE inhibition immediately prior to CS test trials is sufficient to produce extinction retrieval deficits and cause the relapse of extinguished fear.

### Sinusoidal 8 Hz stimulation of the RE prevents fear renewal

Given that RE inactivation impairs the retrieval of extinction memories, we next sought to determine if RE stimulation could enhance extinction retrieval and prevent the relapse of extinguished fear that normally occurs when an extinguished CS is presented outside the extinction context (i.e., renewal)^12,42,43^. For this purpose, we chose to optogenetically stimulate the RE using 8-Hz sinusoidal stimulation, which we hypothesized would mimic the oscillatory synchrony in the mPFC-HPC circuit that is associated with extinction retrieval. To test this, rats were injected with an AAV encoding either an excitatory, blue light activated opsin (AAV8-CaMKII-ChR2-GFP) or a control fluorescent protein (AAV8-CaMKII-mCherry), along with an optic fiber, targeting the RE four weeks prior to behavior (Fig. 6A). Because RE is critical for mPFC-HPC theta synchrony, we chose to use an 8-Hz sinusoidal stimulation pattern during each CS presentation (laser onset 5-sec pre-CS and offset 5-sec post-CS) (Fig. 6B). All animals underwent fear conditioning and extinction as previously described; however, retrieval testing took place in a novel context to test the renewal of extinguished fear (Fig. 6C).

**Figure 6.**
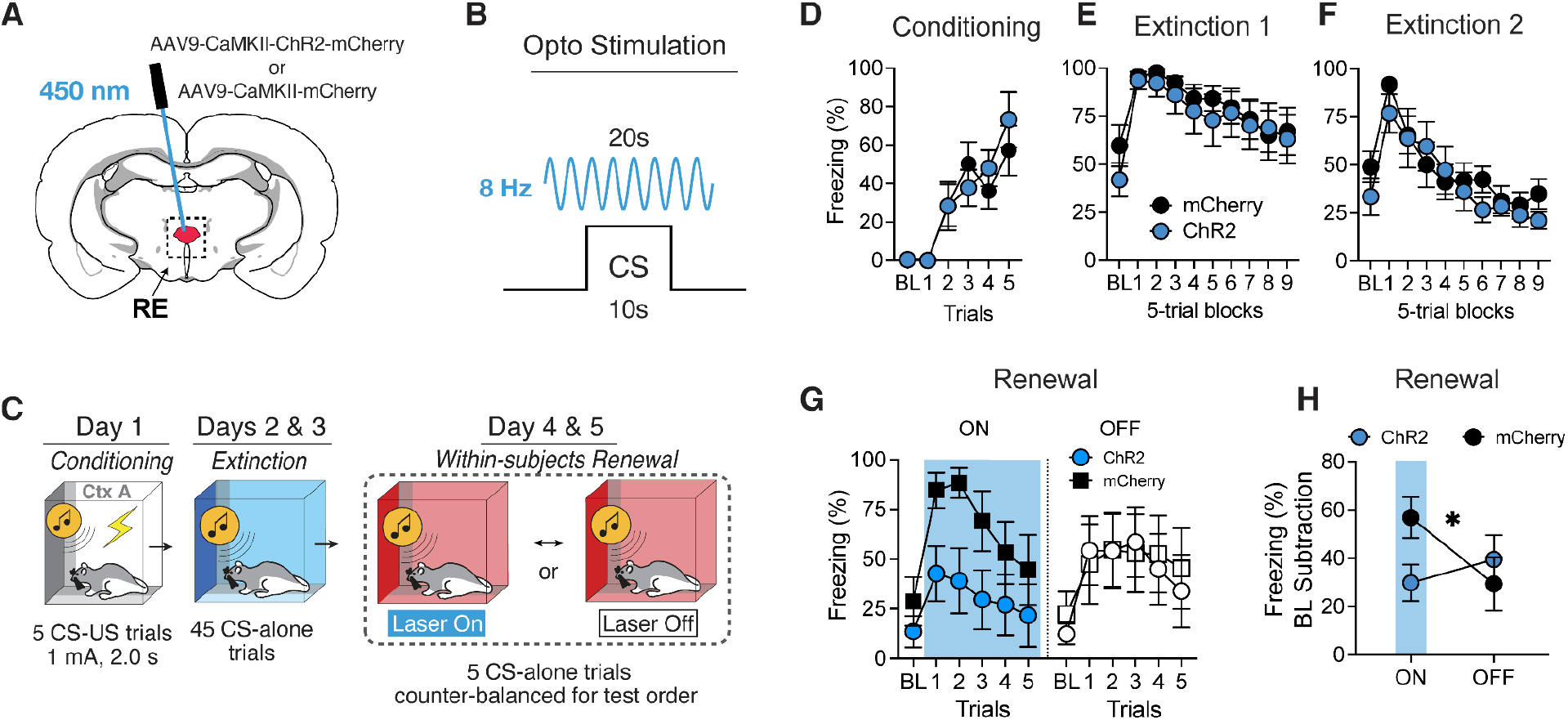
Sinusoidal theta-paced stimulation of the RE reduces the renewal of extinguished fear. (A) Illustration depicting fiber placements and intracranial injection of viruses encoding either the active ChR2 opsin (AAV9-CaMKII-ChR2-GFP) or control mCherry protein (AAV9-CaMKII-mCherry). (B) Schematic of the optogenetic stimulation protocol by which RE neurons were stimulated with an 8 Hz sine wave pattern beginning five seconds before CS onset and stopping five seconds after CS termination. (C) Experimental timeline for fear renewal procedure. Average freezing data during conditioning (D) and extinction sessions (E-F). Average freezing behavior during fear renewal showing that sinusoidal 8 Hz stimulation of RE reduces fear renewal in rats expressing ChR2 (*n* = 6) and not mCherry (*n* = 6) (G-H). All data are means ± s.e.m.s; \**p* < 0.05; \*\**p <* 0.01.

During behavioral testing, rats expressing the light sensitive opsin (ChR2; *n* = 8) and control rats (mCherry; *n* = 8) acquired similar levels of conditioned freezing (Fig. 6E; main effect of Trials: *F*_*5, 70*_ = 18.90, *p* < .0001; Trials x Group interaction: *F*_*5, 70*_ = 0.733, *p*= .601) and extinguished freezing to the CS to similar degrees (Fig. 6F; main effect of Blocks: *F*_*5, 126*_ = 9.081, *p* < .0001; Blocks x Group interaction: *F*_*9, 126*_ = 0.386, *p*= .940). If optogenetic stimulation of the RE blocks fear renewal, one would expect the ChR2 group to display lower fear relative to mCherry controls during the blue light stimulation. Indeed, we found that mCherry rats display high levels of freezing particularly during the first two trials of renewal testing, whereas ChR2 animals remained at low levels of fear (Fig. 6G). This was confirmed by a three-way ANOVA (Trial x Laser x Group interaction: *F*_*5, 65*_ = 2.795, *p* = .024). We also computed a baseline subtraction of the retrieval data to confirm that these effects were not due to elevated levels of background contextual fear in the control rats. Indeed, we found that this effect held even when accounting for differences in basal fear levels between groups (Fig. 6H; Laser x Group interaction: *F*_*1, 13*_ = 5.63, *p* = .034). Thus, theta-paced optostimulation of the RE facilitated extinction retrieval and attenuated the renewal of extinguished fear.

## Discussion

Understanding the neural mechanisms that mediate fear suppression is critical to improving therapeutics for anxiety- and trauma-related disorders^2,14^. Previous work found that the RE and its prefrontal afferents are necessary for the retrieval of fear extinction memories^33,34,44^, however, the mechanism by which the RE mediates this fear suppression remains unknown. Here, we show that 1) fear extinction is characterized by enhanced mPFC-HPC theta synchrony and theta-range oscillations in the RE during fear expression and extinction. Inactivation of the RE impaired mPFC-HPC network activity (c-Fos and theta synchrony) underlying extinction memory retrieval and theta-paced stimulation of the RE rescued context-dependent fear renewal. Collectively, this work shows that the RE mediates fear memory suppression by synchronizing mPFC-HPC theta oscillations.

Substantial work has now demonstrated that limbic brain regions display dissociable 4-Hz and 8-Hz theta-range oscillations during freezing and non-freezing behavior, respectively^16,22,23,45–47^. The slower 4-Hz rhythms arise from respiratory patterns that act to entrain the olfactory bulb, and downstream mPFC and amygdala, to maintain high fear states^45,48–50^. In line with this, we found that increases in 4-Hz power in both the mPFC and RE strongly correlated with increases in freezing behavior throughout fear extinction and retrieval. Interestingly, despite the prominence of 4-Hz rhythms in the mPFC and RE, we found that mPFC-HPC synchrony during fear retrieval and extinction was specific to 8-Hz theta oscillations. This is in line with higher frequency “type-I” hippocampal theta commonly observed in the dorsal HPC during movement, whereas lower frequency “type-II” hippocampal theta is more commonly associated with the ventral HPC (vHPC) during immobility^51–53^. Indeed, the vHPC has been shown to play a more prominent role in processing motivationally-relevant information^54–56^ and selectively synchronizes with the mPFC and amygdala during negatively valenced behaviors^57–61^. Interestingly, it was recently shown that the vHPC synchronizes with and leads the mPFC during fear memory retrieval^62^, the opposite directionality that might be predicted during extinction retrieval^18,33^. Moreover, direct projections from the vHPC provide feedforward inhibition of mPFC neurons to drive fear renewal^12^, a relapse phenomenon not dependent on the RE^33^. Collectively, this provides support for the idea that bidirectional HPC➝mPFC and mPFC➝RE➝HPC interactions act to guide context-appropriate recall of episodic memories^9,63,64^, and that these interactions may be facilitated by neural synchrony in different frequency bands^40,65–67^.

Fear relapse associated with RE inactivation is specifically due to the impaired ability of rats to successfully retrieve “safe” extinction memories. We and others have previously shown that inactivating the RE prior to extinction retrieval causes fear relapse^33,34,44^. However, it was recently found that RE inactivation may be anxiogenic^41^, which might account for increases in freezing behavior after RE inactivation. However, we found that increased freezing after RE inactivation was only exhibited by animals that underwent conditioning and extinction; RE inactivation did not increase freezing in non-conditioned animals. Moreover, pharmacological inactivation of the RE does not increase freezing during baseline periods prior to CS presentations^33^. Although we did not observe differences in freezing during the baseline period, RE activity during the baseline period may be critical for extinction retrieval. Previous work has found that the RE is necessary for properly discriminating contexts^35^, and reductions in context discrimination might result in a relapse of fear^42^. Nonetheless, we found that optogenetic inactivation of RE during CS trials was still sufficient to drive fear relapse. Moreover, we found that 8-Hz sinusoidal stimulation of the RE was sufficient to drive extinction retrieval in a novel context, which normally drives fear renewal. Collectively, this work demonstrates that fear relapse induced by RE inactivation is specific to memory-retrieval deficits and suggests that the RE mediates successful extinction retrieval, not via recognition/encoding of the extinction context, but rather via a CS-related mechanism.

Bidirectional prefrontal-hippocampal interactions are critical for several mnemonic functions, including the encoding and recall of episodic memories^9^. It has been hypothesized that prefrontal top-down control of hippocampal activity via the RE is critical for the suppression of context-inappropriate memories^8^. After extinction, the reduction of conditional freezing behavior in the extinction context requires that animals suppress retrieval of fear memories— memories that normally show strong generalization to any context in which an aversive CS is encountered^4^. Previous work has shown that theta oscillations in the mPFC lead those in the HPC during extinction retrieval^18^ and we have shown that the RE and its afferent projections from the mPFC are critical for extinction encoding and retrieval^33^. The present work reveals that RE coordinates oscillatory synchrony in the mPFC and HPC and that RE inhibition reduces mPFC-HPC coherence and attenuates extinction retrieval. Moreover, theta-paced stimulation of RE can restore extinction retrieval and attenuate fear relapse. These data suggest that mPFC-HPC interactions mediated by the RE are essential to the retrieval operations that permit the suppression of context-inappropriate fear memories in the extinction context. Given that the HPC has also been shown to encode contextual extinction memories^68,69^, it is also conceivable that this circuit is involved in arbitrating the competition between fear and extinction memories for expression in behavior. Given that the RE exerts a predominantly inhibitory influence over the HPC^70^, we speculate that the RE acts to suppress the retrieval of hippocampal fear engrams, thereby facilitating the retrieval of hippocampal extinction engrams.

In summary, the experiments reported here demonstrate that the retrieval of fear extinction memories requires mPFC-HPC interactions that are coordinated by the RE. This work expands our neurobiological understanding of the role of the prefrontal-hippocampal interactions in fear suppression and we suggest that the RE may be a potential therapeutic target for memory-based disorders, such as PTSD.

## Materials and Methods

### Animals

Adult naïve male and female Long-Evans Blue Spruce rats (200 - 240 g upon arrival) were obtained from Envigo (Indianapolis, IN). Rats were individually housed in clear plastic cages in a climate-controlled vivarium with a fixed 14:10 hour light:dark cycle (lights on at 7:00 AM) and all experiments were conducted during the light phase. Rats were given access to standard rodent chow and water ad libitum. Upon arrival, all rats were handled by the experimenter (∼30 sec/rat/day) for a minimum of 5 days prior to the start of any surgical or behavioral procedures. All experimental procedures were conducted in accordance with the US National Institutes of Health (NIH) Guide for the Care and Use of Laboratory Animals and were approved by the Texas A&M University Institutional Animals Care and Use Committee (IACUC).

### Viruses

AAV9-CaMKII-hChR2(H134R)-mCherry, AAV9-CaMKII-mCherry, AAV8-CaMKII-Jaws-KGC-GFP-ER2, and AAV8-CaMKII-GFP viruses were purchased from Addgene (Watertown, MA). All viruses were diluted to a final titer of ∼4-6 × 10^12^ GC/mL with sterilized 1x DPBS.

### Surgery

For all surgeries, animals were anesthetized with isoflurane (5% for induction, 1-2% for maintenance) and placed into a stereotaxic frame (Kopf Instruments). The hair on the scalp was shaved in a separate area, povidine-iodine was applied to the skin and cleaned three times, ocular ointment was applied, and a scalp was used to male a small incision and expose the top of the skull. Finally, the skull was leveled by placing bregma and lambda in the same horizontal plane and small holes were drilled in the skull using round drill burrs for intracranial implants (electrodes. cannulae, and optical fibers) and four small Jeweler’s screws.

For optogenetic experiments, rats received bilateral infusions (0.5 μl) of viruses into the RE. Viruses were infused at a rate of 0.1 μL/min and injector tips were left in the brain for a total of ten minutes to allow for diffusion. The coordinates for RE viral injections were: AP: -2.10 mm, ML: -1.25 mm, DV: -7.09 mm at a 10-degree angle (relative to bregma skull surface). Immediately after viral injection, optical fibers with white ceramic ferrules were implanted into the RE 0.3 mm above the viral injection site [200-μm core; 10-mm length; Thorlabs (Newton, NJ)].

For electrophysiological recordings, a 16-channel microelectrode array (Innovative Neurophysiology, Durham, NC) was chronically implanted targeting the mPFC and the dorsal CA1 region of the HPC unilaterally in the right hemisphere. Microelectrodes targeting the mPFC were comprised of 16 tungsten wires in a 2×8 array such that one row of eight electrodes were 10.5-mm long and the other row was 5-mm long to allow for simultaneous targeting of the infralimbic and prelimbic cortices, respectively. For the HPC, 5-mm long 4×4 arrays were used. All wires were spaced 200-μm from adjacent wires and each wire was 50-μm in diameter. The ground and reference channels were joined together and a single, silver ground wire was wrapped around a skull screw above the cerebellum and affixed with conductive silver paint. For all arrays, the AP coordinate was based on the center of the array and the ML coordinate was based on the leftmost wire of the array. Dental cement was used to secure optical fibers and microelectrode arrays to the skull.

Topical antibiotic (Triple Antibiotic Plus; G&W Laboratories) was applied to the surgical site and one subcutaneous injection of carprofen (5mg/kg) was provided for post-operative pain management. Rats were given a minimum of one week to recover prior to the beginning of behavioral testing.

### Behavioral procedures

Fear conditioning and extinction was conducted in two distinct rooms within the laboratory. Each chamber consisted of two aluminum sidewalls and a Plexiglas ceiling and rear wall, a hinged Plexiglas door, and a grid floor. The grid floor consisted of 19 stainless steel rods that were wired to a shock source and solid-state grid scrambler for delivery of the footshocks (Med Associates). A speaker for delivering auditory stimuli, ventilation fans, and house lights were installed in each chamber. Fear retrieval sessions were conducted in similar chambers (30 × 24 × 21 cm; Med-Associates) equipped with either a red laser (Dragon Lasers, Changqun, China) or electrophysiology system (Plexon, Dallas, TX).

For all conditioning sessions and optogenetic experiments, locomotor activity was acquired online by a load-cell system positioned underneath the behavioral chamber which converted chamber displacements into voltages via Threshold Activity software (Med-Associates). Freezing was defined as less than 10 a.u. for at least one second. For electrophysiological experiments, the load-cell activity was recorded directly via the electrophysiological recording software (OmniPlex, Plexon) to allow for synchronous movement and electrophysiological recordings. For all systems, freezing was defined as immobility that lasted at least 1 s. Therefore, all freezing behavior was automatically recorded and analyzed, providing unbiased measurements.

Distinct contextual environments (contexts A, B, and C) were used during conditioning and retrieval. For context A, the house light was turned off and the overhead white light and ventilation fan were turned off. The cabinet door remained open for the duration of each session. The chamber was wiped with 1.0 % ammonium hydroxide prior to each behavioral session. Rats were transported to context A in black plastic boxes. For context B, the house light was turned on, the fan was turned on, and the room was dimly lit by overhead fluorescent red lights. The cabinet door remained closed for the duration of each behavioral session. A black Plexiglas floor was placed over the grid (except during fear conditioning) and the chamber was wiped down with a 3.0 % acetic acid solution prior to each behavioral session. Rats were transported to context B in white plastic boxes with a clean layer of bedding. For context C, both the house light and overhead white light were turned on, the fan were turned off, and the cabinet door remained closed. Black and white striped wallpapers were taped on the chamber walls. A clear plastic floor was placed over the grid. The chamber was wiped with 70% ethanol prior to each behavioral session and rats were transported to context C in white plastic boxes with a clean layer of bedding.

Distinct contexts were always used during conditioning and retrieval. Context A was composed of white ambient room lighting, house light turned off, cabinet doors remained open, a metal grid floor, and conditioning chambers were cleaned with 1.0% ammonium hydroxide. For context B, red ambient room lights were turned on, house lights were turned on, cabinet doors were closed, a smooth black plastic floor, and chambers were cleaned with 3.0% acetic acid. Finally, for context C, both the house light and white room light were turned on, one chamber door remained open, black and white striped wallpapers were taped onto the exterior of clear chamber walls, a textured white plastic floor, and the chambers were cleaned with 70% ethanol.

For all behavioral experiments, rats underwent a standard auditory fear conditioning procedure in which an innocuous auditory tone (conditioned stimulus; CS) was paired with an aversive footshock (unconditioned stimulus; US) in context A. This conditioning session was comprised of a 3-min baseline period and five CS (10 s, 80 dB, 8 kHz)-US (1.0 mA, 2 s) pairings with 60-s intertrial intervals (ITIs), and additional 60-s post-shock period. Unless otherwise noted, all rats underwent fear extinction twenty-four hours later in context B. This consisted of a 3-min baseline period followed by 45 CS-alone presentations (10s, 80 dB, 30s ITI). Some animals that showed resistance to fear extinction underwent multiple rounds of extinction until adequately low levels of fear were observed. For optogenetic and pharmacological testing, rats underwent a within-subject retrieval session such that animals were tested under Light/Drug conditions in a counterbalanced manner.

### Drug infusions

For drug microinfusions, rats were moved into a room adjacent to the vivarium and placed into 5-gallon white buckets. Dummy cannula internals were removed and a stainless-steel injector (33-gauge, 9 mm; Plastics One) connected to polyethylene tubing was inserted into the guide cannula. Polyethylene tubing was connected to 10-μl Hamilton syringes that were mounted in an infusion pump (Kd Scientific). Muscimol was diluted to a concentration of 0.1 μg/μl in sterile saline. Infusions were made at a rate of 0.3 μl/min for 1 min and the injectors were left in place for 2 min post-infusion to allow for adequate diffusion. Each infusion was verified by movement of an air bubble that separated the drug or sterile saline from distilled water within the polyethylene tubing. Clean dummy internals were inserted into each guide cannula after infusions. All infusions were made ∼5 min prior to behavioral testing. All animals were acclimated to these procedures by performing dummy changes twice before beginning behavioral procedures.

### Optogenetics

Rats expressing Jaws or control GFP in RE neurons were illuminated using a red laser (635 nm laser; Dragon Lasers, Changqun, China). Rats expressing ChR2 and their controls were illuminated using a blue laser (450 nm laser; Dragon Lasers, Changqun, China). All optic fibers were hand constructed and only fibers with >80% efficiency after polishing were used. Laser power was set to 10 mW at the end of the patchcord for all experiments to ensure ∼8-9 mW at the tips of optical fibers. A fiber-optic rotary joint (Doric Lenses Inc; Québec, QC, Canada) and a bundled patch cord (Thorlabs; Newton, NJ) were used to allow optostimulation while not impeding locomotion or exploratory behaviors. For “constant” inhibition experiments, the laser was controlled by Med Associates software via a TTL adaptor and light was delivered 10 s before the first CS onset and persisted until the end of the testing session. A DG812 Waveform Generator (RIGOL Technologies, Inc.) driven by TTL riggers was used to generate 8 Hz sinusoidal stimulation patterns.

### Electrophysiology

Local field potentials (LFPs) and freezing behavior were automatically recorded by a multichannel neurophysiological recording system (OmniPlex, Plexon, Dallas, TX). All electrophysiological data were acquired at 40k Hz and then down-sampled to 1000 Hz and saved for offline analysis of local field potentials.

Local field potential analyses were conducted using a combination of custom written MATLAB and Python scripts. All data were detrended and notch filtered at 60 Hz (58-62 Hz window) to remove power line noise. The raw data for all trials were plotted and any trials with noticeable motion artifacts were manually removed from all analyses. For power analyses, power spectral density (PSD) estimates were calculated using Welch’s method (*pwelch* MATLAB function). PSDs were then averaged across trials to obtain one PSD estimate per subject for each time point of interest. Average power in frequency bands of interest (3-6 Hz and 6-9 Hz) were then averaged to obtain a single frequency-band power estimate for each subject. For coherence analyses, the magnitude squared coherence was also calculated using Welch’s method (*mscohere* Matlab function) and averaged in the same way as power calculations. All spectrograms and coherograms were created using the multitaper method from the Chronux toolbox using a 3 s moving window with 100 ms overlap.

### Histology

After the end of behavioral experimentation, rats were sacrificed to confirm electrode, cannula, viral, and/or fiber optic placement. To help visualize electrode tip placements, a small current (0.1 mA) was passed through the four of the sixteen microelectrodes for 10 s to create a small lesion immediately before transcardial perfusions. Animals were overdosed with sodium pentobarbital (Fatal Plus, 100 mg/ml, 0.7 ml), transcardially perfused with ice-cold saline and fixed with 10% physiological formalin.

Perfused brains were placed in physiological formalin for 14–24 h before being moved to a 30% sucrose solution for a minimum of three days. After three days, or until brains had sunk, all brains were frozen and sectioned at −20° on a cryostat. To verify cannula and electrode placements, brains were sectioned at 40 μm, mounted onto gelatin subbed slides, thionin stained (0.25%), cover-slipped with Permount (Fisher Scientific), and then imaged on a wide-field stereoscope. To confirm viral expression and optic fiber localization, brains were sectioned at 30 μm, mounted onto gelatin subbed slides, cover-slipped with DAPI-infused Fluoromount (Fisher Scientific), and then imaged on a 10x fluorescent microscope.

### Immunohistochemistry

All rats were overdosed with sodium pentobarbital and transcardially perfused. Brains were dissected and stored in 10% formalin for up to 14 hours and then transferred to 30% sucrose at 4°C for at least 72 hours. After all brains were sectioned on a cryostat (−20°) at 30 μm and brain sections. For Fos immunohistochemistry, brain sections were washed 3x in TBST, incubated in 0.3% H20h for 15 min, washed 3x in TBS, and then incubated in rabbit anti-c-fos primary antibody (1:1000; Millipore) overnight. The next day, The sectioned were washed 3x in TBS followed by a one-hour incubation in a biotinylated goat anti-rabbit secondary antibody (1:1000, Jackson Immunoresearch), amplified with the avidin biotin complex at 1:1000 (ABC; Vector labs), and visualized with 3,3’-diaminobenzidine (DAB) + nickel ammonium sulfate. Stained sections were mounted in subbed slides, coverslipped with Permount and stored at room temperature until they were imaged.

To quantify c-Fos expression, one 10X image (895 μm × 670 μm; 0.596 mm^2^) was taken of each hemisphere at different AP levels of each the PL and IL (+2.7mm and +2.3mm from bregma), HPC (-5.5mm and -6.0mm), and MGN (-5.5mm and -6.0mm). The total number of Fos-expression neurons in each image was manually counted, averaged across all images for each image, and divided by the surface area to the average number of Fos cells/mm^2^.

### Statistics

All data were analyzed using custom-written Matlab and Python scripts. Statistical analyses of the data were performed using GraphPad Prism (version 9.0; GraphPad Software). There were no sex differences in any of the analyses; males and females were collapsed. Data were submitted to repeated-measures analysis of variance, and significant interactions were followed by Tukey’s multiple comparisons test. Group sizes were determined based on prior work. All data are represented as mean ± SEM, and p < .05 was considered statistically significant.

## Acknowledgements

We thank Dr. Jianfeng Liu for his assistance with the optogenetic experiments. We also thank Drs. Justin Moscarello, Rachel Smith, and Ursula Winzer-Serhan for helpful discussions around the direction and interpretation of this work. This work was supported by the National Institutes of Health (R01MH065961 and R01MH117852 to S.M.).

## Author contributions

MT and SM designed all electrophysiological and optogenetic experiments; KR, JJ, and SM designed the c-Fos experiment. KR and JJ performed c-Fos immunization and counting. MT and SP performed optogenetic experiments; MT analyzed the data. MT performed all other experiments and analyzed the data. MT and SM wrote the manuscript.

## Data availability

The data from these experiments are available from the corresponding author upon request.

